# Lifespan reference curves for harmonizing multi-site regional brain white matter metrics from diffusion MRI

**DOI:** 10.1101/2024.02.22.581646

**Authors:** Alyssa H. Zhu, Talia M. Nir, Shayan Javid, Julio E. Villalon-Reina, Amanda L. Rodrigue, Lachlan T. Strike, Greig I. de Zubicaray, Katie L. McMahon, Margaret J. Wright, Sarah E. Medland, John Blangero, David C. Glahn, Peter Kochunov, Asta K. Håberg, Paul M. Thompson, Neda Jahanshad, Alzheimer’s Disease Neuroimaging Initiative

**Affiliations:** Imaging Genetics Center, USC Mark and Mary Stevens Neuroimaging and Informatics Institute, Keck School of Medicine of USC, Marina del Rey, CA, USA; Department of Biomedical Engineering, USC Viterbi School of Engineering, Los Angeles, CA, USA; Department of Psychiatry and Behavioral Science, Boston Children’s Hospital, Harvard Medical School, Boston, MA, USA; QIMR Berghofer Medical Research Institute, Brisbane, QLD, Australia; Queensland University of Technology, Brisbane, QLD, Australia; Queensland Brain Institute, University of Queensland, Brisbane, QLD, Australia; Centre for Advanced Imaging, University of Queensland, Brisbane, QLD, Australia; School of Psychology, Brisbane, QLD, Australia; Department of Human Genetics, University of Texas Rio Grande Valley, Brownsville, TX, USA; South Texas Diabetes and Obesity Institute, University of Texas Rio Grande Valley, Brownsville, TX, USA; Olin Neuropsychiatry Research Center, Institute of Living, Hartford, CT, USA; Faillace Department of Psychiatry and Behavioral Sciences at McGovern Medical School, The University of Texas Health Science Center at Houston, Houston, TX, USA; Department of Neuromedicine and Movement Science, Faculty of Medicine and Health Sciences, Norwegian University of Science and Technology (NTNU), Trondheim, Norway; Department of MiDtT National Research Center, St. Olav’s Hospital, Trondheim University Hospital, Trondheim, Norway

## Abstract

Age-related white matter (WM) microstructure maturation and decline occur throughout the human lifespan, complementing the process of gray matter development and degeneration. Here, we create normative lifespan reference curves for global and regional WM microstructure by harmonizing diffusion MRI (dMRI)-derived data from ten public datasets (N = 40,898 subjects; age: 3-95 years; 47.6% male). We tested three harmonization methods on regional diffusion tensor imaging (DTI) based fractional anisotropy (FA), a metric of WM microstructure, extracted using the ENIGMA-DTI pipeline. ComBat-GAM harmonization provided multi-study trajectories most consistent with known WM maturation peaks. Lifespan FA reference curves were validated with test-retest data and used to assess the effect of the ApoE4 risk factor for dementia in WM across the lifespan. We found significant associations between ApoE4 and FA in WM regions associated with neurodegenerative disease even in healthy individuals across the lifespan, with regional age-by-genotype interactions. Our lifespan reference curves and tools to harmonize new dMRI data to the curves are publicly available as eHarmonize (https://github.com/ahzhu/eharmonize).

## Introduction

Methodological variability and small sample sizes have contributed to poor reproducibility in population studies across many research fields, including neuroscience and brain imaging^1,2^. As brain MRI scanning is costly and time-intensive, large well-powered neuroimaging studies typically require pooling of data from smaller, existing studies or data collection across multiple sites. In either case, two primary sources of variance make it non-trivial to combine multi-site neuroimaging data. First, heterogeneous study design and subject inclusion criteria may yield results that are generalizable only to similar cohorts. Second, MRI scans and derived metrics vary considerably due to differences in scanner hardware and software, acquisition parameters such as spatial or angular resolution, and image processing pipelines^3^. Even when acquisition protocols are harmonized, inter-site differences still persist^4^.

Multi-study consortia, such as the Enhancing NeuroImaging Genetics through Meta-Analysis (ENIGMA) consortium^5^, may account for study differences using a traditional two-stage (i.e., group-based) meta-analysis. Meta-analyses allow individual studies to account for study-specific covariates and population substructure, but if specific traits or conditions of interest have a low prevalence, e.g., rare copy number variants in the genome, meta-analysis may not be possible as single sites may lack sufficient samples to run statistical models. When small studies are conducted, model assumptions of normally distributed sampling and known variances may be violated^6^, and inflated effect sizes from smaller studies may contribute to the meta-analytic results^1^. Meta-analyses may also be unable to distinguish between site differences and confounding biological variables that are correlated with site^7^. Beyond this, in a lifespan study, age-dependent effects (e.g., specifically occurring in development or senescence) may be diluted using a meta-analysis approach.

Brain imaging studies that analyze data from individuals across the lifespan may benefit instead from a “mega”-analysis approach, which pools data or derived imaging features for a single analysis based on all individual data points. Mega-analysis can accommodate a broader range of statistical models and better control of effect size estimation^8^. Growth charts have been established for anthropometric measurements, such as height and weight, but until recently, study differences and the lack of standardization have hindered pooling datasets of sufficiently large sample sizes and age ranges to establish normative models for brain MRI measures. The emergence of large public datasets and data harmonization techniques have made lifespan brain charts possible to create. Initial ‘brain charts’ using structural MRI measures across the lifespan have combined multi-study data for mega-analyses with different approaches to account for site heterogeneity, including generalized additive models for location, scale, and shape (GAM-LSS) to account for non-linear trajectories of global and regional volume, surface area, and thickness measurements^9^, linear and non-linear hierarchical Bayes models of subcortical volume and cortical thickness^10,11^, and fractional polynomial regression applied to harmonized cortical thickness measures^12^. Ge et al. compared methods for normative modeling of brain morphometry and chose multivariate fractional polynomial regression models to create sex-specific lifespan charts for subcortical volumes and cortical surface area and thickness measures, available through CentileBrain (www.centilebrain.org)^13^.

ComBat is a commonly used harmonization method, initially developed to correct for batch effects in gene expression arrays^14^, but later applied to derived neuroimaging measures^15,16^. ComBat expanded upon previous models, implementing an empirical Bayes framework that is robust to small sample sizes. Recent extensions of ComBat include ComBat-GAM^17^ and CovBat^18^, which account for non-linear effects of covariates on the brain measures, and adjust for site differences in data covariance, respectively. Structural imaging studies have shown that ComBat and ComBat-GAM may improve the detection of case-control group differences in schizophrenia and post-traumatic stress disorder (PTSD) cohorts compared to unharmonized mixed-effects models that model study as a random effect^19,20^.

The initial application of ComBat in the neuroimaging literature was to data derived from diffusion-weighted MRI^16^. Diffusion-weighted MRI (dMRI) is sensitive to white matter (WM) microstructure and is influenced by a greater number of acquisition parameters than T1-weighted MRI, including spatial and angular resolution, diffusion times, *b*-value, and the number of shells^21^. In general, greater angular resolution and number of *b*-shells (e.g., diffusion weightings) give more stable diffusion metrics^22,23^, and smaller voxel sizes are less susceptible to partial voluming but may have more noise ^22,24^. The initial application of ComBat to dMRI-derived data was limited to pooling together only two studies that had similar age ranges and acquisition parameters^16^. While it has since been used more widely in studies of epilepsy^25^, traumatic brain injury^26^, and neurogenetic disorders^27^, harmonization of dMRI metrics across age ranges and acquisition protocols has not been explored in depth. Single-site dMRI studies have found lifespan trajectories of WM microstructure to be different to those of T1w-derived gray matter measures: peak WM maturation typically occurs over a decade later than corresponding peaks for gray matter metrics^28,29^, necessitating the development of dMRI-specific lifespan curves. Most prior dMRI studies have used measures extracted from the diffusion tensor imaging (DTI) model. The DTI model has been widely adopted in clinical studies to understand WM microstructure because, compared to more advanced models, it can be reliably fit to lower resolution data which is more convenient and faster to collect. This widespread use is particularly advantageous when combining studies with different dMRI acquisitions.

Although harmonization approaches such as ComBat are widely used, limitations remain. ComBat regularizes site-specific harmonization parameters differently depending on sample size, and is sensitive to the input data, such that if a new cohort were to join the study, the harmonization would need to be repeated. To avoid incorrectly modeling true biological variability as a scanner-related effect, datasets must have a sufficient overlap in all covariates, e.g., age ranges. Multi-site mega-analyses often need a centralized database, which may limit participation by sites that cannot share individual level data. Harmonizing data to a template or reference that can be distributed would overcome this limitation. ComBat and its variants typically adjust input data so that the residuals of a statistical model have the same mean and variance, but none provide a reference dataset to perform normalization consistently across studies. However, care is required when choosing the reference dataset, as harmonized results may be biased toward the selected data. An ideal reference for harmonization would include multiple populations across a wide age range, such as the NiChart Reference Dataset aggregating structural MRI data for the iSTAGING consortium^17^, and provide a means to account for differences in MRI acquisition protocols. A normative model could be created, incorporating heterogeneous datasets to represent the different sources of variance, and used as a reference for ComBat harmonization. Here, we created a framework that harmonizes input data to built-in lifespan reference curves, allowing for distributed harmonization, easy incorporation of new datasets, and version control of updatable lifespan reference curves.

Here, we harmonized DTI data from ten public datasets, totaling 13,297 subjects (aged 3-95), to create lifespan reference curves for white matter regions extracted with the ENIGMA-DTI pipeline^30,31^. The ENIGMA-DTI pipeline is a well-established pipeline based on tract-based spatial statistics^32^ with standardized outputs, which has been run on data from over a hundred cohorts worldwide, in studies of over ten disorders^33^. We evaluated lifespan curves computed using ComBat, ComBat-GAM, and CovBat to determine which methods would most effectively capture the expected non-linear trends^28,34,29,28,34^. After creating an optimal set of lifespan reference curves, we harmonized an independent group of test datasets (2,161 subjects, aged 3-85), acquired with dMRI protocols previously unseen in our training data, to our newly established lifespan reference curves. We characterized the impact of acquisition parameters on harmonization parameters, the regional sex differences in our lifespan reference curves, and the performance of our framework on longitudinal data. Lastly, as the effect of genetic risk factors may vary across the lifespan, we evaluated our framework by assessing the effects of Apolipoprotein E (ApoE) genotype on white matter microstructure in a multi-study sample aged 3-85 (N = 30,915). The E4 allele is the major risk haplotype for late-onset Alzheimer’s disease - compared to the most common genotype, E3 - whereas E2 is less common and may have a neuroprotective effect^35^. The overall framework - including the harmonization mechanism and our lifespan reference curves are freely available in an open source Python package called ENIGMA Harmonize (eHarmonize; https://github.com/ahzhu/eharmonize).

## Methods

### Datasets

#### Data for Building Reference Curves

Public dMRI datasets collected from individuals across a variety of age ranges were combined to span 3 to 95 years of age (N = 40,898 subjects; 47.6% male; Table 1). The pediatric and adolescent cohorts (age: 3-21) included the Pediatric Imaging, Neurocognition, and Genetics (PING) dataset^36^; Human Connectome Project in Development (HCP-D)^37^; and the Adolescent Brain and Cognitive Development (ABCD)^38^ studies. As the ABCD study contains twin and sibling data, one sibling per family was randomly selected to create an unrelated subset. Participants in the ABCD study were recruited from a very narrow age range (9-10 years old), yet at the time of analysis, two-year follow up scans had also been released for most participants. We broadened the age range by including data from half the participants at baseline and an independent half at two year follow-up. No additional exclusion criteria were applied to the PING and HCP-D datasets for this work.

**Table 1.** Study demographics and image acquisition parameters of each dataset. In multi-shell acquisitions, the shells used for DTI processing are indicated in **bold**. * Original acquisition dimensions. GE automatically zero-pads in *k*-space by default, so processing was performed on resampled data. ^‡^ QTIM participants were scanned twice, once with each protocol. ^‡‡^ Non-zero *b*-value shells were processed separately, for comparison.

| Study/Dataset | Exclusion Criteria | Age Range (years) | Available N (%) Male | Bootstrapped N (%) Male | Country | Manufacturer (Field Strength) | $b$-values (volumes) | Resolution (mm<sup>3</sup>) |
| --- | --- | --- | --- | --- | --- | --- | --- | --- |
| PING | - | 3-21 | 906 (52.2) | 902 (52.1) | United States | Siemens (3T)<br>GE (3T)<br>Philips (3T) | **0 (4), 1000 (32)**<br>**0 (2), 1000 (30)**<br>**0 (3), 1000 (30)** | 2.5 x 2.5 x 2.5<br>2.5 x 2.5 x 2.5*<br>2.5 x 2.5 x 2.5 |
| HCP-D | - | 8-21 | 617 (48.5) | 617 (48.5) | United States | Siemens (3T) | **0 (28), 1500 (93), 3000 (92)** | 1.5 x 1.5 x 1.5 |
| ABCD | One sibling per family | 9-13 | 5611 (52.7) | 5000 (52.8) | United States | Siemens (3T) | **0 (7), 500 (6), 1000 (15), 2000 (15), 3000 (60)** | 1.7 x 1.7 x 1.7 |
| SLIM | - | 17-26 | 432 (45.1) | 432 (45.1) | China | Siemens (3T) | **0 (1), 1000 (30)** | 2.0 x 2.0 x 2.0 |
| HCP | One sibling per family | 22-36 | 363 (45.5) | 363 (45.5) | United States | Siemens (3T) | **0 (18), 1000 (90), 2000 (90), 3000 (90)** | 1.25 x 1.25 x 1.25 |
| Cam-CAN | - | 18-87 | 305 (35.1) | 305 (35.1) | United Kingdom | Siemens (3T) | **0 (3), 1000 (30), 2000 (30)** | 2.0 x 2.0 x 2.0 |
| PPMI | Parkinson's disease | 30-81 | 83 (63.9) | 83 (63.9) | International | Siemens (3T) | **0 (1), 1000 (64)** | 1.98 x 1.98 x 2.0 |
| UK Biobank | ICD-10 codes: C70-C72, G, I6, Q0, R90, R940, S01-S09; self-report conditions : 1081, 1086, 1266, 1267, 1397, 1434, 1437, 1491, 1626 | 45-82 | 31,986 (46.8) | 5000 (47.0) | United Kingdom | Siemens (3T) | **0 (8), 1000 (50), 2000 (50)** | 2.0 x 2.0 x 2.0 |
| OASIS3 | Cognitive impairment | 42-92 | 225 (48.4) | 225 (48.4) | United States | Siemens (3T) | **0 (1), 1000 (64)** | 2.5 x 2.5 x 2.5 |
| ADNI3 | Cognitive impairment | 55-95 | 370 (41.2) | 370 (41.2) | United States<br>Canada | Siemens (3T)<br>Siemens (3T)<br>GE (3T)<br>GE (3T)<br>Philips (3T)<br>Philips (3T) | **0 (1), 1000 (30)**<br>**0 (7), 1000 (48)**<br>**0 (4), 1000 (32)**<br>**0 (6), 1000 (48)**<br>**0 (1), 1000 (32)**<br>**0 (4), 1000 (32)** | 2.0 x 2.0 x 2.0<br>2.0 x 2.0 x 2.0<br>2.0 x 2.0 x 2.0*<br>2.0 x 2.0 x 2.0*<br>2.0 x 2.0 x 2.0<br>2.0 x 2.0 x 2.0 |
| NIH Pediatric | Age < 3 years | 3-21 | 122 (44.3) | - | United States | Siemens (1.5T)<br>GE (1.5T) | **0 (10), 100 (10), 300 (10), 500 (10), 800 (30), 1100 (50)**<br>**0 (9), 100 (10), 300 (10), 500 (10), 800 (30), 1100 (50)** | 2.5 x 2.5 x 2.5<br>2.5 x 2.5 x 2.5 |
| TAOS | One sibling per family | 12-16 | 232 (49.6) | - | United States | Siemens (3T) | **0 (3), 700 (55)** | 1.7 x 1.7 x 3 |
| GOBS | Kinship > 0.125 | 23-78 | 189 (37.0) | - | United States | Siemens (3T) | **0 (3), 700 (55)** | 1.7 x 1.7 x 3 |
| QTIM-d30<sup>†</sup> | One twin per family | 18-29 | 317 (35.6) | - | Australia | Bruker (4T) | **0 (3), 1159 (27)** | 1.8 x 1.8 x 5.5 |
| QTIM-d105<sup>†</sup> | One twin per family | 18-29 | 317 (35.6) | - | Australia | Bruker (4T) | **0 (11), 1159 (94)** | 1.8 x 1.8 x 2 |
| HUNT | Multiple sclerosis, cysts, meningioma, infarction, post-op changes | 50-67 | 811 (46.7) | - | Norway | GE (1.5T) | **0 (5), 1000 (40)** | 2.5 x 2.5 x 2.5* |
| ADNI2 | Cognitive impairment | 60-85 | 42 (40.5) | - | United States Canada | GE (3T) | **0 (5), 1000 (41)** | 2.7 x 2.7 x 2.7 |
| ADNI3-S127<sup>††</sup> | Cognitive impairment | 61-85 | 64 (34.4) | - | United States | Siemens (3T) | **0 (13), 1000 (48), 2000 (60)** | 2 x 2 x 2 |

The young adult cohorts (age: 17-36) included baseline data from the Southwest University Longitudinal Imaging Multimodal (SLIM) brain data repository^39,40^ and Human Connectome Project (HCP)^41^. Each study acquired MRI data on a single scanner. As with ABCD, HCP is family-based, so only one sibling per family was selected. The Cam-CAN dataset included younger, midlife, and older adults (18 to 87 years)^42^. Mid-adulthood to older adults (age: 30-95) were included from Tier 1 of the Parkinson’s Progression Markers Initiative (PPMI)^43^, Open Access Series of Imaging Studies (OASIS3) dataset^44^, the population-based UK Biobank study^45^, and a subset of the third phase of the Alzheimer’s Disease Neuroimaging Initiative with single-shell diffusion imaging acquisition (ADNI3)^46^. In the PPMI, OASIS3, and ADNI3 studies, participants with a diagnosis of Parkinson’s disease, mild cognitive impairment, or dementia were excluded from the reference training cohort. Exclusion criteria applied to the UK Biobank included neurological and cerebrovascular disorders as well as incidental findings on brain MRI and head injuries.

#### Evaluation Datasets

Seven datasets were held out to evaluate the reference curves made from the other datasets, as detailed in the **Reference Curve Evaluation** section (Table 1).

Children scanned as part of the NIH Pediatric MRI study with the extended diffusion MRI protocol were included as the pediatric test cohort^47^. Children younger than 3 years of age were excluded. Brain MRIs from the Teen Alcohol Outcomes Study (TAOS) and the Genetics of Brain Structure and Function (GOBS) study were acquired on the same scanner with the same acquisition protocol^48^. As a result, they were combined to provide a lifespan dataset including both children and adults. One child per family was selected for the TAOS dataset. As the GOBS study is composed of extensive pedigrees, selecting one member per family would have decreased the sample size considerably. Instead, a kinship filter was implemented, where only individuals with less than a first-degree cousin relationship were included (kinship < 0.125). Older adults from the Trøndelag Health Study (HUNT) study were also included^49,50^.

To further evaluate our template in handling test data with differences in scan parameters, we included two datasets with which we calculated DTI metrics for the same individuals in different ways. The first dataset, the Queensland Twin IMaging (QTIM) study^51^, included individuals scanned with two protocols of differing angular and spatial resolution^52^. The second dataset we used was the subset of multi-shell diffusion MRI data from ADNI3, for which we calculated DTI metrics with both *b*-shells.

### Image processing

A summary of dMRI acquisition parameters for template building and evaluation datasets can be found in Table 1. In the training datasets with a multi-shell acquisition, the most appropriate shell for diffusion tensor imaging (DTI) was chosen for processing, usually *b* = 1000 s/mm^2 53,54^. As with many large-scale multi-study initiatives in ENIGMA, preprocessing steps varied across studies as datasets were collected and processed over the past 15 years. Preprocessing followed guidelines provided by the ENIGMA-DTI team (on GitHub), which includes steps such as EPI distortion and eddy current distortion correction. FSL’s *dtifit* was used to calculate fractional anisotropy (FA) and diffusivity maps. As ADNI3-S127 dMRI acquisitions were multi-shell, *dtifit* was run on volumes from the *b*=1000 and *b*=2000 s/mm^2^ shells independently, for comparison.

Images were then processed through the ENIGMA-DTI pipeline^31^. The FA maps were warped to the ENIGMA template and then skeletonized using tract-based spatial statistics^32^. The mean FA value was extracted from the full skeleton as well as each region of interest (ROI) defined by the JHU atlas^30^. Bilateral ROIs were combined by averaging the measures across hemispheres, and some subregion ROIs were combined to create a single measure for the entire structure, e.g., the three parts of the corpus callosum. In both cases, the combined average was weighted by the number of voxels in each ROI. A lifespan reference curve was created for all measures, lateralized and combined. In subsequent analyses, we report only the results for the combined measures (25 ROIs; Table 2).

**Table 2.**
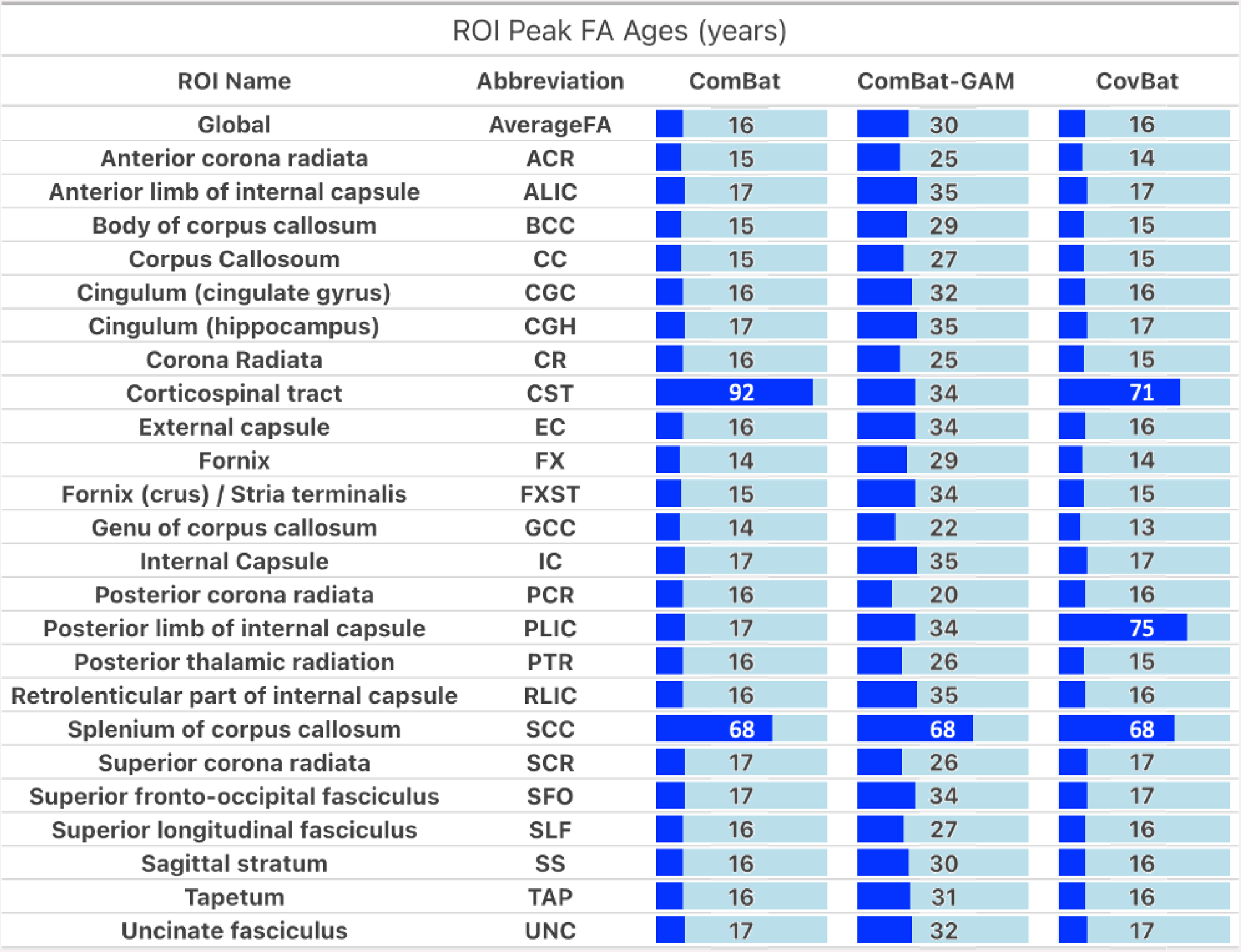
The age at which regional white matter skeleton FA trajectories peaked fell mostly below 20 years of age in the ComBat and CovBat reference curves. In the ComBat-GAM reference, the peak ages largely fell between the ages of 20 and 40 years old, which Kochunov and Lebel had both previously reported. One exception is the SCC which had a peak age of 68 across methods. The CST from the JHU atlas has shown low reliability, so the results for this ROI should be interpreted with caution^72^. The *tapetum* and uncinate fasciculus reported here reflect the current JHU atlas labels rather than the ENIGMA-DTI lookup table, which was based on a previous JHU atlas version.

| ROI Peak FA Ages (years) |  |  |  |  |
| --- | --- | --- | --- | --- |
| ROI Name | Abbreviation | ComBat | ComBat-GAM | CovBat |
| **Global** | **AverageFA** | 16 | 30 | 16 |
| Anterior corona radiata | ACR | 15 | 25 | 14 |
| Anterior limb of internal capsule | ALIC | 17 | 35 | 17 |
| Body of corpus callosum | BCC | 15 | 29 | 15 |
| Corpus Callosum | CC | 15 | 27 | 15 |
| Cingulum (cingulate gyrus) | CGC | 16 | 32 | 16 |
| Cingulum (hippocampus) | CGH | 17 | 35 | 17 |
| Corona Radiata | CR | 16 | 25 | 15 |
| Corticospinal tract | CST | 92 | 34 | 71 |
| External capsule | EC | 16 | 34 | 16 |
| Fornix | FX | 14 | 29 | 14 |
| Fornix (crus) / Stria terminalis | FXST | 15 | 34 | 15 |
| Genu of corpus callosum | GCC | 14 | 22 | 13 |
| Internal Capsule | IC | 17 | 35 | 17 |
| Posterior corona radiata | PCR | 16 | 20 | 16 |
| Posterior limb of internal capsule | PLIC | 17 | 34 | 75 |
| Posterior thalamic radiation | PTR | 16 | 26 | 15 |
| Retrolenticular part of internal capsule | RLIC | 16 | 35 | 16 |
| Splenium of corpus callosum | SCC | 68 | 68 | 68 |
| Superior corona radiata | SCR | 17 | 26 | 17 |
| Superior fronto-occipital fasciculus | SFO | 17 | 34 | 17 |
| Superior longitudinal fasciculus | SLF | 16 | 27 | 16 |
| Sagittal stratum | SS | 16 | 30 | 16 |
| Tapetum | TAP | 16 | 31 | 16 |
| Uncinate fasciculus | UNC | 17 | 32 | 17 |

### Quality Control

We limited the ABCD subset in our training sample to include only data from Siemens scanners, after preliminary quality assurance as detailed in Supplementary Figure 1. In the PPMI dataset, where controls could have prodromal Parkinson’s disease, dMRI data underwent visual quality control (QC), and those with movement artifacts were excluded. Statistical QC was also performed by removing subjects who had any ROI metric fall outside of 5 standard deviations of the site mean.

**Figure 1.**
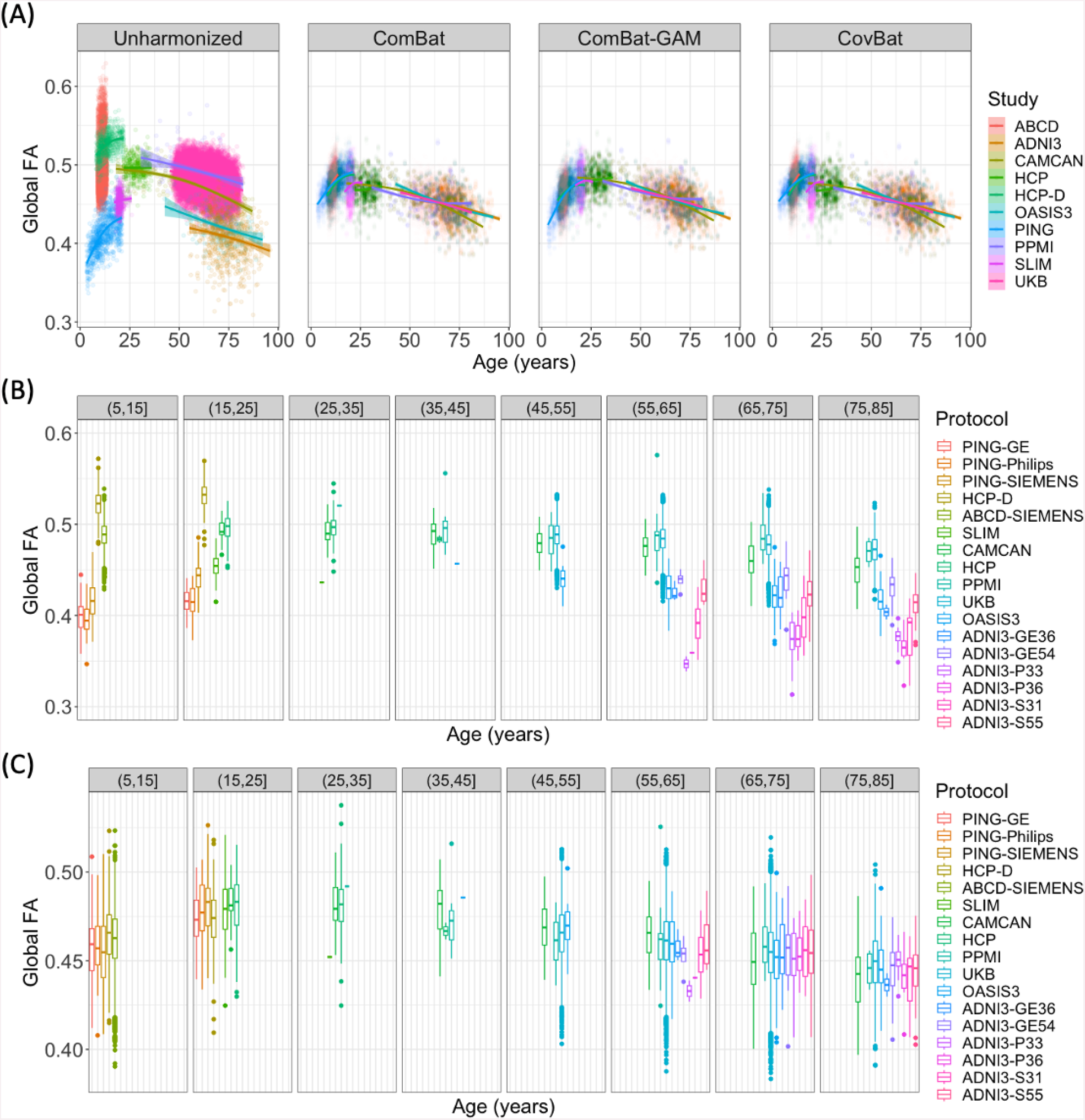
**(A)** After harmonization using different methods, average FA in the full WM skeleton is plotted against age and colored by study. Age-binned boxplots of **(B)** unharmonized data and **(C)** data harmonized using ComBat-GAM show the median global FA were quite different between protocols pre-harmonization and were more similar post-harmonization.

### Harmonization Methods

Regional FA metrics from the training datasets were harmonized using three approaches: ComBat, ComBat-GAM, and CovBat ^14,17,18^. ComBat and ComBat-GAM were run using the *neuroHarmonize* package in Python (https://github.com/rpomponio/neuroHarmonize). CovBat was run using the R package of the same name (https://github.com/andy1764/CovBat_Harmonization). Across all methods, age and sex were used as covariates, with the dataset modeled as the batch effect. As the outputs of the ComBat family harmonization methods are weighted by the sample size of each study, harmonization was run using iterative subsampling of all datasets to provide more equal weighting across datasets (25 iterations; 200 subjects per dataset where available; 13,297 subjects in total) as the UK Biobank greatly outnumbered the other datasets. The larger datasets (UK Biobank and ABCD) were sampled without replacement, while the smaller datasets were sampled with replacement.

#### ComBat^14^

ComBat is a location and scale adjustment model where, for site *i* and individual *j*, it assumes that the feature measurements are modeled as a linear combination of site effects and non-site effects, written as:

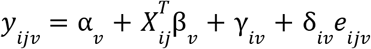

where *α*_*v*_ is the overall mean per feature *v*, *β*_*v*_ is the vector of corresponding coefficients to the covariate matrix *X, γ*_*iv*_ is an offset from the grand mean per site *i* and feature *v, e*_*ijv*_ is the residual vector, and *δ*_*iv*_ is the multiplicative site effect of site *i* on feature *v*. ComBat removes the additive and multiplicative effects from the residuals using

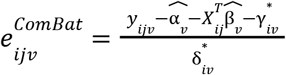

where 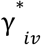 and 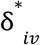 are estimated using an empirical Bayes framework. The harmonized data are then calculated by

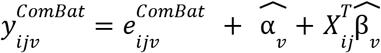

where 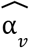 and 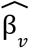 are the respective parameter estimates.

#### ComBat-GAM^17^

ComBat-GAM extends ComBat by using generalized additive models (GAM^55^) to model non-linear covariate effects. Modeling of the non-linear covariates is achieved by placing one or more covariates within the function, f(x):

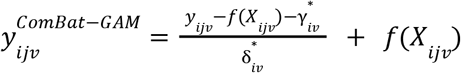

where *f*(*X*_*ijv*_) is a smooth function over the covariates^56^.

#### CovBat^18^

CovBat was proposed to harmonize both mean and covariance batch effects in data for multivariate pattern analysis. First, CovBat applies ComBat to normalize the mean and variance of the residuals in a statistical model with for *p* features. The ComBat-adjusted residuals, 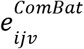, are calculated for each feature independently, but may still retain site-specific covariance. Thus in the second step, principal component analysis is performed on the residuals of the full dataset to identify covariance patterns and reduce the number of dimensions. The ComBat-adjusted residuals can now be expressed as

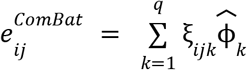

where 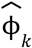are the estimated principal components obtained as the eigenvectors of the full-data covariance matrix, *ξ*_*ijk*_ are the principal component scores, and *q* is the number of orthogonal axes.

Treating the principal component scores analogously to those of the original features, site-specific covariance can be removed via

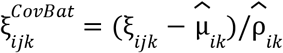

where 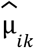 and 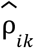 are the site-specific center and scale parameters, respectively, of each principal component.

Finally, the CovBat-adjusted residuals are obtained by projecting the adjusted scores into residual space via

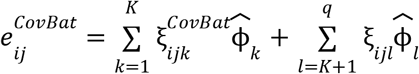

where *K* is the number of principal components chosen to capture the user-specified percent variation.

Then the final CovBat-adjusted observations are obtained by adding the intercepts and covariates estimated in the first step using ComBat:

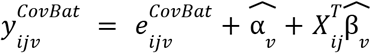

### Creating the Reference Curves

After the training datasets were subsampled and harmonized using the three ComBat methods, the outputs of each method were combined for evaluation. To create reference curves, GAM models were fit for each ROI and harmonization method, covarying for sex and smoothing across age using the *mgcv* package in R(basis function: 10 cubic splines). To evaluate the harmonization methods, we extracted the peak age, i.e., the age of maximum FA, from each region that exhibited a concave down trajectory. We compared our harmonized curves to previous single-site studies of the white matter microstructure that reported FA to peak between the ages of 20 and 40 years old ^28,29^. We used these as our “silver standard”, i.e., a standard established by *in vivo* models rather than histological studies.

Once the optimal harmonization method was selected, GAM models were fit to each ROI, covarying for sex and smoothing across age. We used the *qgam* package in R (basis function: 10 cubic splines) to fit quantile regression GAM models for each centile (0.1-0.99) to generate normative lifespan reference curves for each sex.

The ComBat-GAM Python package (neuroHarmonize) does not allow the user to specify a reference site, so we adapted the code to allow for that functionality (available through neuroHarmonize fork: https://github.com/ahzhu/neuroharmonize). Evaluation datasets were harmonized to the lifespan reference curves, i.e., the location and scale parameters (*γ*^*^_*iv*_ and *δ*^*^_*iv*_ respectively) of each site was calculated in relation to the mean and variance of the lifespan reference curves.

### Reference Curve Evaluation

#### Characterizing the Reference Curves

In addition to age-at-peak comparisons, we also characterized the trajectories of our regional reference curves compared to that of the global FA. We calculated the Fréchet distance (FD), a curve similarity metric, between the global FA reference curve and that of each ROI.

We tested for sex differences in bilaterally averaged ROIs. GAM models were fitted to model sex effects on regional FA while covarying for age as a smoothed term (basis function: ten cubic splines). We also tested for smoothed age-by-sex interactions. Multiple comparisons were corrected for using the false discovery rate method (FDR; 25 tests)^57^.

#### Train vs. Test Datasets

Using the adapted ComBat-GAM code, all available data from the training datasets - as well as the seven held-out datasets - were harmonized to the newly created lifespan reference curves. Using each subject’s age and sex, the differences between reference-predicted and site-harmonized values were calculated, and the mean absolute error (MAE) was calculated for each protocol and ROI. For each ROI, an unpaired *t*-test was used to compare the performance of our harmonization framework on training vs test datasets (FDR-corrected; 25 tests).

#### Acquisition Effects on Model Parameters

Spatial resolution, number of diffusion directions, and choice of *b-*value are all known to affect DTI values. We tested for correlations between voxel volume and the ComBat output scale and shift parameters from both training and test datasets. We also analyzed the angular resolution and number of volumes (*b0* and *b*-shells combined); other acquisition parameters such as *b*-value and scanner manufacturer, were highly homogeneous across studies, and we were not able to determine their effects on model parameters.

#### Longitudinal Studies

To determine how our framework would perform for longitudinal studies, we harmonized longitudinal data in two ways: first, we calculated the scale and shift parameters from the baseline data and applied them to the follow-up data; second, we calculated the scale and shift parameters from the baseline and follow-up data separately. We then ran mixed-effects models to determine how the different methods impacted the modeled age effects on each ROI, covarying for sex, age-by-sex interaction, and age^2^ and including subject ID as a random effect. The mixed-effects models were also run in the unharmonized data, which was used for comparison. We used data from the UK Biobank, which has a subset of subjects with follow-up imaging visits acquired approximately two years after baseline (N = 1,384, baseline ages 47-80 years, mean time interval: 2.25 years).

#### Case Studies

To examine the outcome of case-control analyses after harmonization, we chose to analyze the effects of ApoE across the lifespan as the data was available in most datasets and requires no harmonization of its own. Genetic data was collected in seven of the ten training datasets, and two of the evaluation datasets (Table 3). From datasets that released full genetic data, the ApoE SNPs rs7412 and rs429358 were extracted for analysis. Other datasets focusing on aging populations had only made ApoE genotypes available, and the provided data was used for this study. As genome-wide data was not available for all studies, subjects were filtered for European ancestry using self-provided race or ethnicity information. Only healthy controls as defined in Table 1 were included. Analyses were run separately for E2 and E4 allele counts, each using E3E3 homozygotes as controls and excluding carriers of the other allele. Linear regressions tested for effects of E2 or E4 count on harmonized regional FA, adjusting for age, sex, age-by-sex, and age^2^, and multiple comparisons corrected using FDR (25 tests). In secondary analyses, ApoE-by-age interactions were tested in nominally significant ROIs.

**Table 3.**
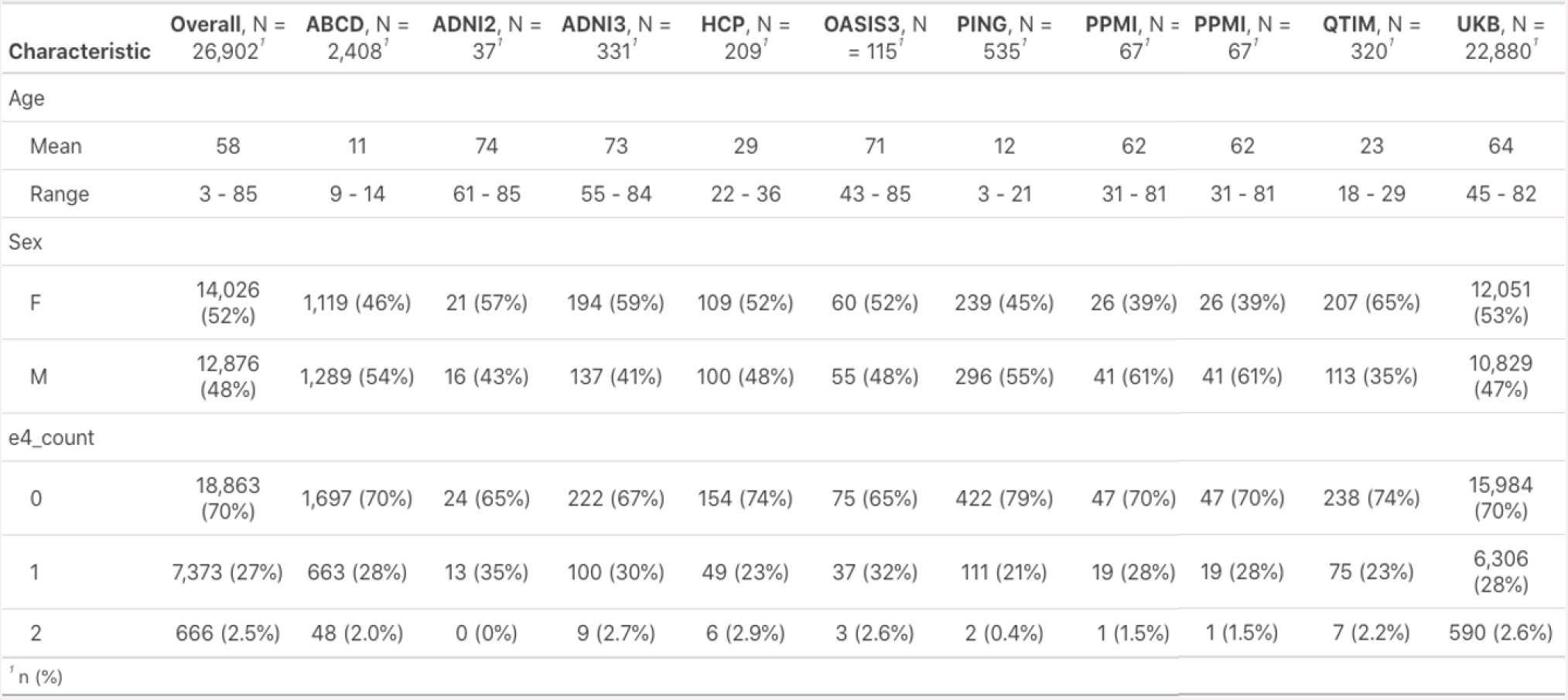
Demographics and ApoE4 information for the datasets included in the regression models. The included subjects are healthy controls filtered for European ancestry.

Two test sets ADNI3-S127 and QTIM had data available from different dMRI protocols on the same subjects, as described above. For both datasets, we ran the ApoE regressions pre- and post-harmonization with both sets of FA metrics. We then compared the effect sizes to determine how much of an impact the differences in protocol made on the statistical outputs, and if they were different, to determine if harmonization would result in a convergence of the results. In the QTIM study, we ran the ApoE4 analyses in both the low spatial and angular resolution scans and the high spatial and angular resolution scans. In the ADNI3-S127 dataset, we ran the same models in the different diffusion shells: *b*=1000 vs *b*=2000 s/mm^2^.

### eHarmonize

Combining our modified ComBat-GAM code with the lifespan reference curves, we created the eHarmonize Python package (https://github.com/ahzhu/eharmonize), which comes equipped with command line tools to read in FA measures and harmonize them to the included centile reference curves while taking age and sex into account (Figure 6). The eHarmonize command line interface was written to harmonize data from a new site to the built-in lifespan reference curves and apply an existing harmonization model to new data from a known site. To account for dMRI acquisitions that do not cover a full field-of-view, eHarmonize detects which subset of ROIs are provided before calculating harmonization model parameters as the underlying neuroHarmonize does not handle missing data.

## Results

### Harmonization methods

The lifespan full WM skeleton FA references that were created are shown in Figure 1. Qualitatively, study FA trajectories across age were better aligned after harmonization, regardless of method. The performance of ComBat and CovBat appeared similar. The peak ages for white matter FA in these references were mostly before twenty years of age (Table 2). In the pediatric datasets, the steep increase in FA with age was maintained, but due to the larger age range and number of adult datasets, the linear model resulted in larger harmonized FA values than the outputs from the GAM model. The peak age of white matter FA in almost all of the ComBat-GAM references was between 20 and 40 years old, matching the expected values ^28,29^. As a result, we used the ComBat-GAM harmonized data to create the lifespan reference curves.

### The Lifespan Reference

As our harmonization was conducted with iterative sampling, we were able to plot a prospective reference for each iteration (Figure 2A). The iterations produced largely consistent results, although the sparse sampling of older subjects (age > 85 years old) resulted in larger confidence intervals at the older ages. Using the outputs from all iterations, we created one set of centile reference curves per sex, covering much of the lifespan (3-95 yrs) (Figure 2B).

**Figure 2.**
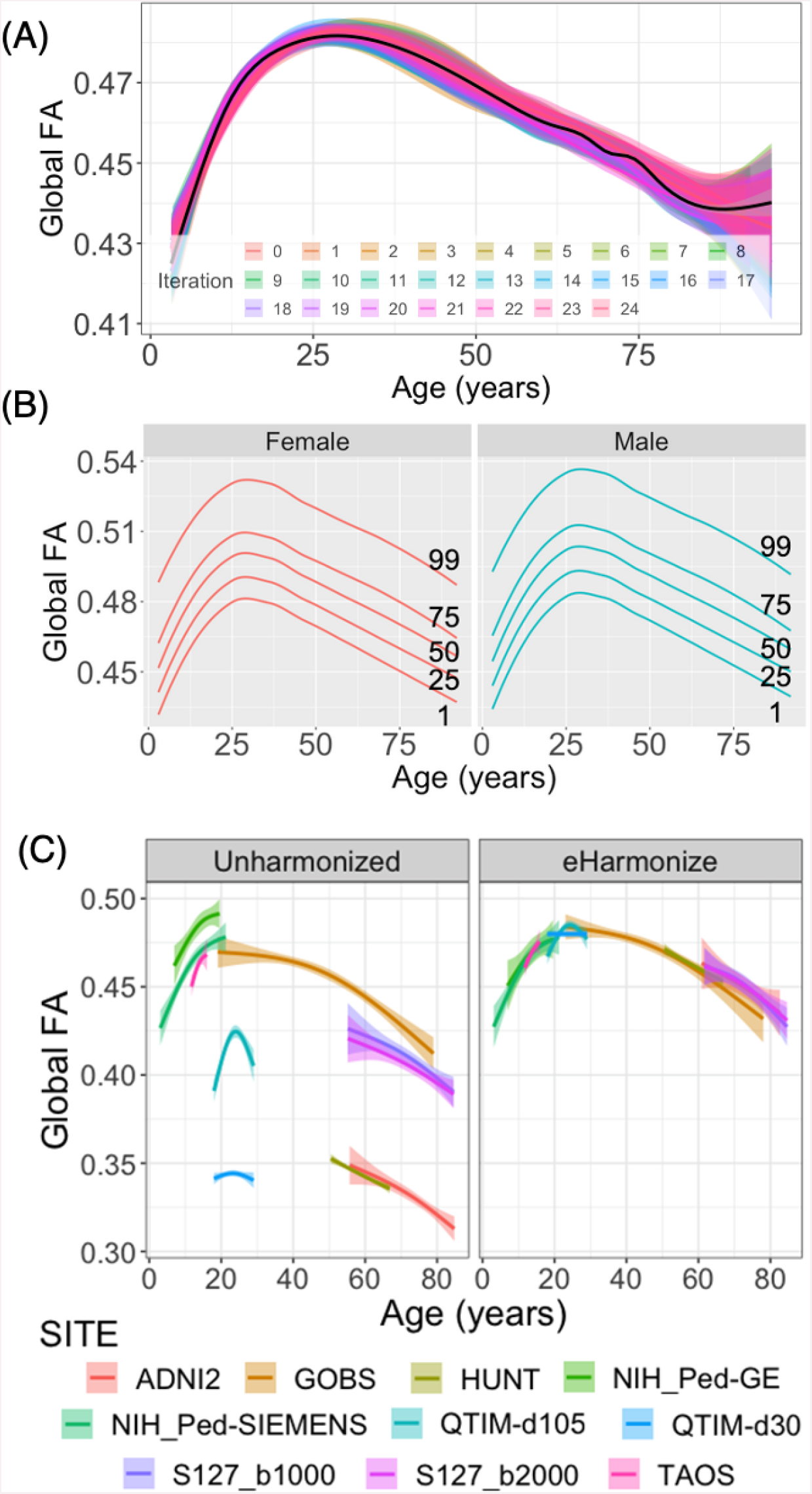
**(A)** Iteration-specific reference curves of the global FA measure as created by iterative subsampling of ∼200 participants from each study and ComBat-GAM harmonization are displayed (25 iterations; mean in black). **(B)** Sex-specific centile curves derived from the results of iterative subsampling harmonization make up the final lifespan reference curve. **(C)** After applying our framework (eHarmonize) to held-out evaluation datasets, the harmonized datasets fall in line with the global FA lifespan reference curve.

#### Characterizing the Reference Curves

With the exception of the splenium of the corpus callosum (SCC), the age peaks of all lifespan reference curves fell between 20 and 40 years of age. Rather than peaking in the 20-40 age range, the SCC plateaus at that age range before rising again for its later age peak. In comparison to the global FA reference curve, most ROIs had lifespan trajectories with a high curve similarity (FD between 0.005 and 0.04). The exception was the fornix (FX), which had a Fréchet distance of 0.14 and a steeper slope of decline after the peak. Lifespan trajectories for SCC and FX can be found in Supplementary Figure 2.

Higher FA was found in females as compared to males in six ROIs: the fornix, the fornix/*stria terminalis*, posterior *corona radiata*, posterior thalamic radiation, sagittal *stratum*, and the *tapetum*. No significant sex differences were found in the body, splenium, or whole corpus callosum. The remaining ROIs showed higher FA in males compared to females. Effect sizes for all FA measures are reported in Supplementary Table 1. We found no significant age-by-sex interactions.

### Post-Harmonization Analyses

#### Train vs. Test Datasets

We harmonized the FA values of all training and test datasets to the newly created lifespan reference curves (Figure 2C). The MAE comparison between training and test datasets found no significant differences across ROIs (p > 0.09).

#### Acquisition Effects on Model Parameters

We extracted the scale and shift parameters for all protocols. Of the tested acquisition parameters, voxel size showed significantly negative correlations with the shift parameter (*γ*\*; -0.57 < r < -0.27) but no correlation with scale (*δ*\*) across most ROIs (Figure 3). Neither the number of directions nor volumes showed a significant impact on either the shift or scale parameters.

**Figure 3.**
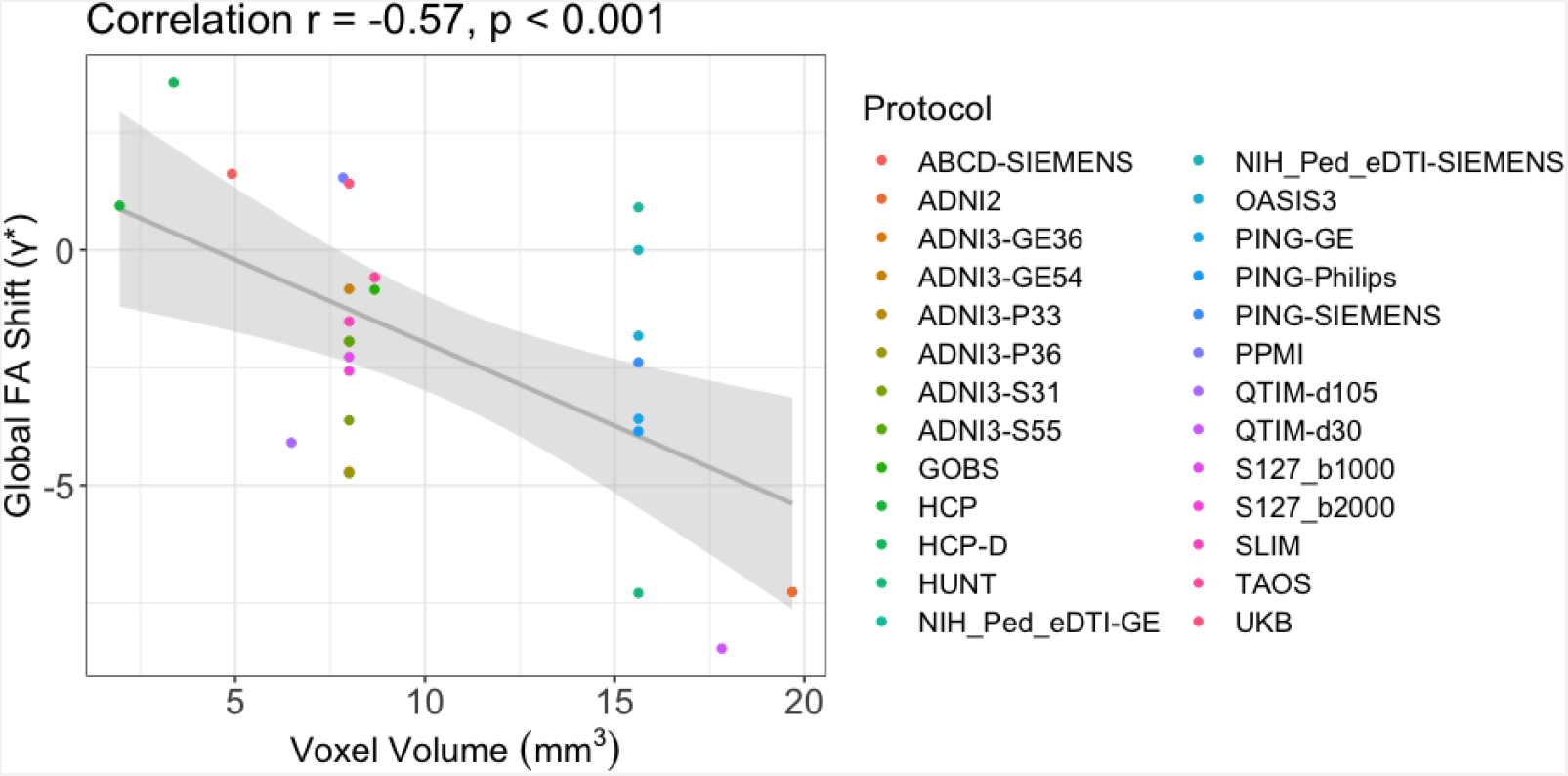
For most ROIs, the shift parameter extracted from the ComBat-GAM model was significantly correlated with voxel volume. The negative correlation between the global FA shift parameter and voxel volume is shown here (*r* = -0.60; p < 0.001).

#### Case Studies

Nine datasets had ApoE data available for analysis. In total, 26,902 subjects (3 to 85 years of age) were included in the E4 analyses (Table 3) and 22,760 aged 3-85 years in E2 analyses. No significant associations were found between E2 count and regional FA. In E4 carriers, significantly lower FA was found in the hippocampal cingulum (CGH; β = -0.027, *p* = 4.1×10^−6^), posterior thalamic radiation (PTR; β = –0.022, *p* = 1.6×10^−4^), overall skeleton (β = -0.020, *p* = 6.2×10^−4^) and splenium of the corpus callosum (SCC; β = -0.016, *p* = 6.8×10^−3^). Effect sizes for all ROIs may be found in Supplementary Table 2. Nominally significant ROIs included the sagittal *stratum*, overall corpus callosum, *genu* of the corpus callosum, retrolenticular part of the internal capsule, and the fornix (crus)/*stria terminalis* (FX/ST). Secondary analyses in all ROIs passing the nominal significance threshold showed a significant age-by-ApoE4 interaction in the FX/ST (β = -7.7×10^−4^; *p* = 0.014).

In datasets with multiple protocols, a comparison of regional ApoE4 standardized beta estimates from regression models run pre-harmonization found similar results between the ADNI3-S127 protocols differing only in *b*-value (r_β_ = 0.97) and less similar results between the QTIM protocols differing in both spatial and angular resolution (r_β_ = 0.60). In the ADNI3-S127 dataset (N = 56), the CGH was found to be nominally significant in both protocols (*b*=1000: β = -0.30, p = 0.016; *b*=2000: β = -0.30, *p* = 0.019), but in the posterior corona radiata, a nominally significant result was only found in the *b*=2000 dataset (*b*=1000: β = -0.26, p = 0.060; *b*=2000: β = -0.28, *p* = 0.046). There were no significant associations in the QTIM dataset (N = 316). After harmonization, the ApoE4 standardized betas of all individual datasets and protocols remained almost identical, ensuring harmonization does not change individual dataset findings (Figure 4).

**Figure 4.**
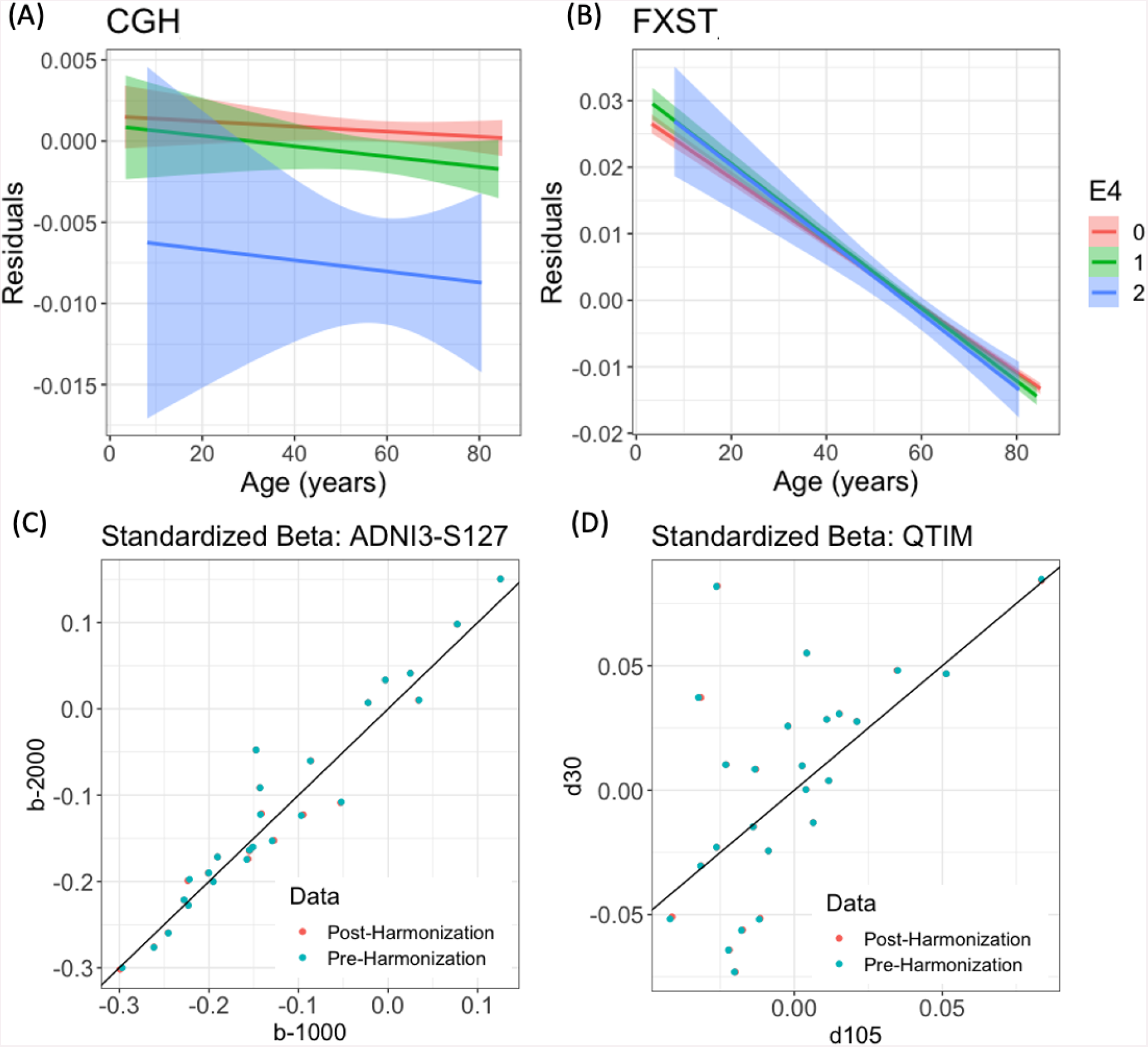
Age associations, residualized by sex, age-by-sex, and age^2^, are plotted for the **(A)** CGH and **(B)** FXST. In the CGH, E4 carriers had significantly lower FA compared to their E3E3 counterparts. In the FXST, the FA of E4 carriers was higher at younger ages but after approximately age 55 years, dropped below that of non-carriers in older ages. A comparison of protocols in the **(C)** ADNI3-S127 and **(D)** QTIM datasets showed that harmonization does not converge the results of the same subjects acquired with different protocols. Each scatter point reflects the association (standardized beta) between ApoE4 and an ROI, corrected for age, sex, age-by-sex, and age^2^.

#### Longitudinal Studies

In the UK Biobank, mixed-effects models detected insignificant differences in associations between age and regional FA from regressions run on raw unharmonized data and data harmonized by baseline parameters; correlations of the age effects between methods, calculated across ROIs, approximated to *r* ∼ 1.0 (Figure 5). When baseline and follow-up data were harmonized independently, the age effect correlation with the raw unharmonized models was 0.92.

**Figure 5.**
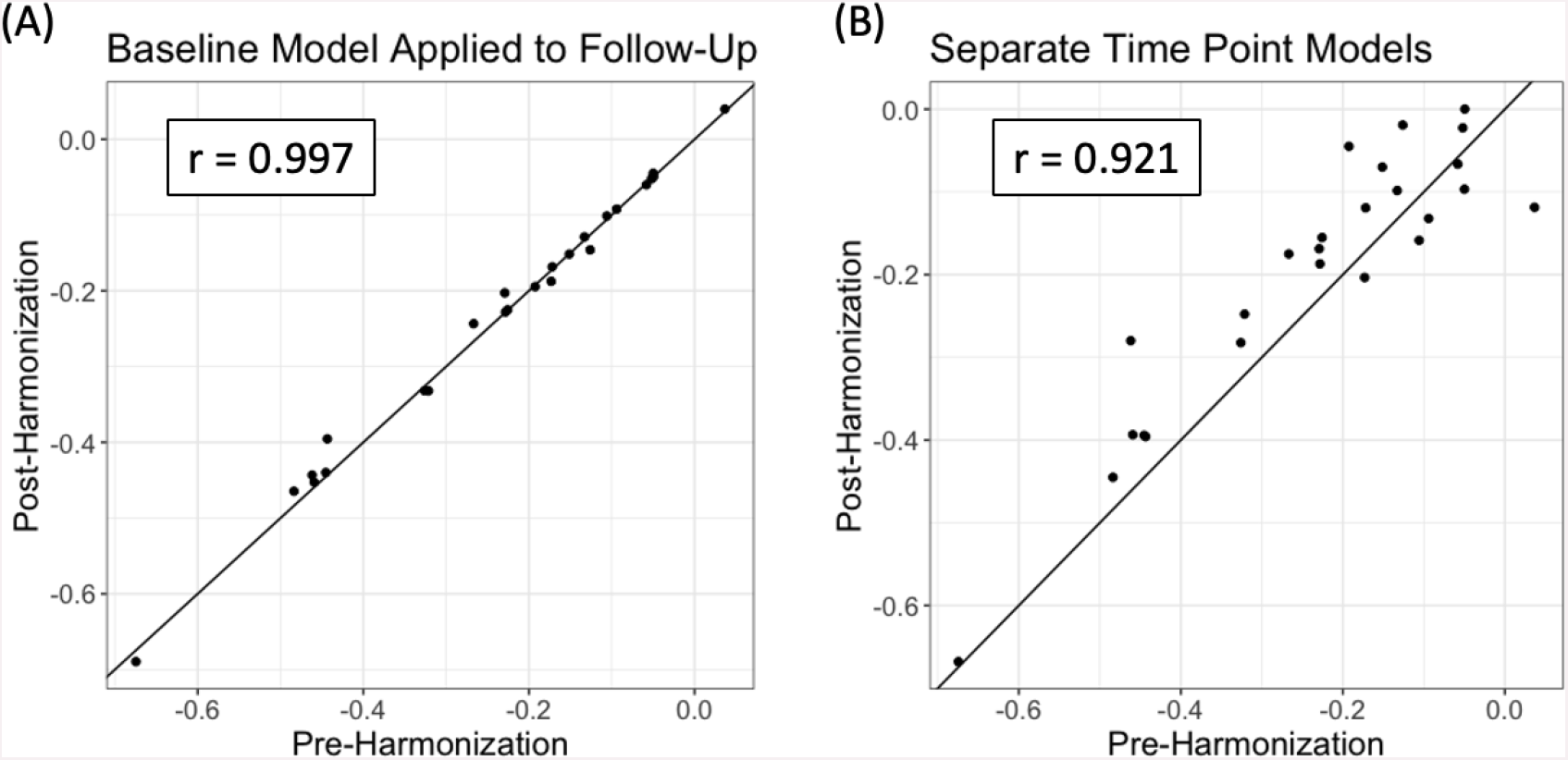
Standardized betas for age associations with each WM ROI either pre-harmonization (x-axis) or post-harmonization (y-axis). Harmonization was either performed by **(A)** applying baseline parameters to follow-up data, or **(B)** modeling the ComBat parameters for each time point separately.

**Figure 6.**
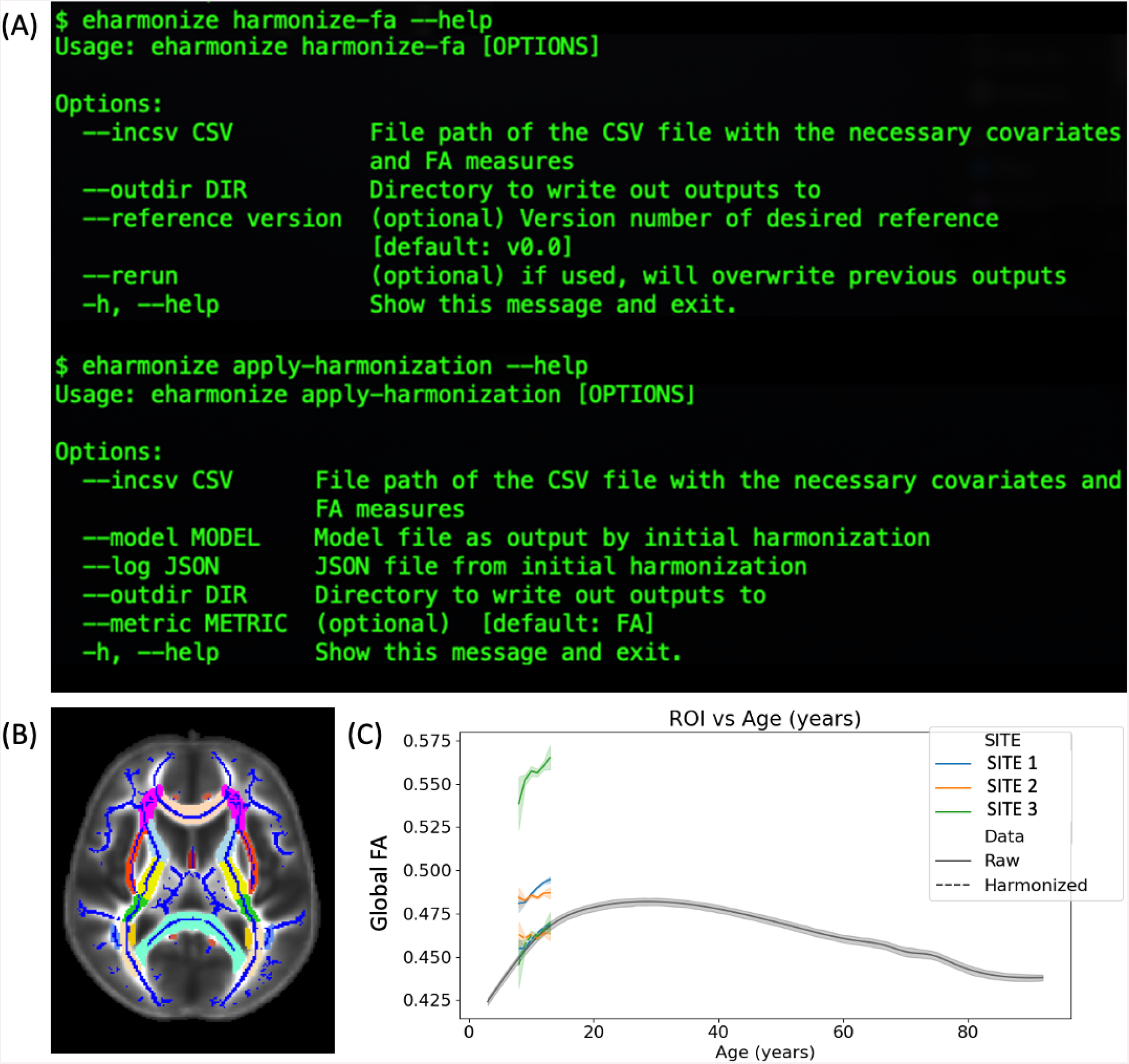
**(A)** The eHarmonize command line interface comes with two subcommands: *harmonize-fa* for harmonizing data from a new site, and *apply-harmonization* for applying an existing harmonization model to a known site. **(B)** The ENIGMA-DTI template with the skeleton and ROIs overlaid. **(C)** QC output showing data before and after harmonization in relation to the reference curve, shown in gray.

### eHarmonize

Our package is set up to be adaptable. In addition to the lifespan reference curves, a JSON file is included containing meta information about the reference curves (e.g., version number, datasets used in their creation). This feature allows for updated references to be implemented in the future while preserving previous versions for ongoing studies to maintain consistency. References for other diffusion measures, or measures from any other modality (imaging or non), can easily be implemented by adding the appropriate information to the meta JSON.

The outputs of eHarmonize include the harmonized data, model parameters, and a text log file for provenance, reflecting a timestamp of when the tool was run and by whom, and the reference and ROIs that were used. If a study includes cases and controls, eHarmonize will harmonize the measures based on the controls and then apply the model to the cases as is done in the neuroHarmonize package. A QC image per ROI is also output showing a line plot of the study data vs. age before and after harmonization, with the reference in the background (Figure 6C).

## Discussion

In this work, we combined dMRI data from ten public datasets to create lifespan reference curves for global and regional white matter (WM) fractional anisotropy (FA). We found that ComBat-GAM best matched the previously reported non-linear age trends across the lifespan and expected age peaks seen in single cohort studies of WM development. Across most regions, our reference curves show a steep increase in FA during development, peaking between the ages of 20 and 40, followed by a continuous and gradual decrease for the rest of the lifespan. One notable exception was the splenium of the corpus callosum. While the FA also increases steeply during development, it plateaus in the early 20s and then peaks later at 68 years. Previous studies have found the FA of the splenium of the corpus callosum (SCC) to be relatively stable with age in adulthood^58^, possibly due to posterior-to-anterior development and anterior-to-posterior aging trends^59,60^. The widely diverging projections of the splenium may account for the late peak as the loss of diverging fibers increases FA followed by an overall decline^61^. Another outlier region is the fornix, which has the same overall trends, but a much sharper decline. Given its location, this likely reflects greater misregistration and partial voluming associated with age-related atrophy and nearby ventricular expansion^62^.

We found sex differences across the WM skeleton across the lifespan. Most regions showed higher FA in males than in females, and the opposite effect was found in six ROIs. Prior studies of sex differences in white matter microstructure have also reported regional variation^50,63,64^. These sex effects may be affected by covariates, such as intracranial volume (ICV). In a study using the HUNT dataset, analyses without an ICV covariate found regionally varying sex effects, but after including ICV, only females had regions with significantly higher FA^50^. We did not incorporate ICV as a covariate in our harmonization framework as it is not an output of the ENIGMA-DTI pipeline and is generally estimated from T1-weighted MRI, as opposed to dMRI.

For longitudinal studies, we showed that harmonization parameters modeled on baseline data can be applied to follow-up data or parameters can be modeled independently in each timepoint. Results from our comparisons to unharmonized data were largely consistent between the two methods. In the UK Biobank, which we used for the analyses, the time interval between visits was much smaller than the age range of the study population. In some studies, the follow-up age range may fall outside the modeled age range at baseline, and a separate follow-up model may be advisable. This may be particularly advantageous for data in the non-linear ranges of the lifespan reference curves.

One major benefit of a lifespan dMRI reference is that it can be used to study subtle effects, such as genetic influences, on brain WM microstructure throughout life. Here, we harmonized FA data from the healthy controls of ten datasets to our lifespan reference curves and found an effect of lower global FA in subjects with the ApoE4 genotype compared to E3E3 homozygotes. The same effect was found regionally in the hippocampal cingulum, the posterior thalamic radiation, and the splenium of the corpus callosum. As a genetic risk factor for Alzheimer’s disease (AD), our findings overlap with regions previously implicated in DTI studies of dementia^65,66^. Previous studies of white matter microstructure in cognitively healthy controls have also found E4 carriers to have lower FA than non-carriers, though published results are inconsistent and some studies have also reported null findings^67^. Most previous studies were either limited by smaller sample sizes (N < 200) or a limited age range with most focusing on individuals aged 60 years and above, and fewer than five studies incorporating younger individuals. Our study was conducted in over 30,000 subjects and across the lifespan.

We also found a significant age-by-E4 interaction in the fornix (crus)/*stria terminalis* whereby the FA of the E4 carriers was higher than that of E3 homozygotes early in life until approximately age 55 years, after which E4 carriers showed lower FA than noncarriers. The fornix and the *stria terminalis* are major output tracts of the hippocampus and amygdala, respectively. In a longitudinal lifespan study of structural imaging measures, age-dependent associations were found between E4 and rates of volume change in the hippocampus and the amygdala^7^. In both structures, E4 was associated with prolonged growth into adulthood and faster atrophy later in life. This age-by-E4 interaction may reflect the ‘antagonistic pleiotropy’ hypothesis that the ApoE4 genotype may be advantageous earlier in life^68^.

As part of our ApoE analyses, we further evaluated datasets that had subjects acquired with multiple dMRI protocols. Diffusion-weighted MRI, and FA in particular, is affected by many acquisition parameters^21^. We found that the use of our lifespan harmonization framework does not converge the statistical results of the same population acquired on different protocols. However, we note that differences in FA due to acquisition protocols may reflect different underlying anatomy or biological processes. We found that voxel size was negatively correlated with the harmonization shift parameter, i.e., sites with larger voxels generally have lower FA values. Larger voxels are more likely to contain crossing fibers, which would result in lower FA values. In such a case, convergence of FA values between protocols at the expense of biological interpretability may not be desired.

We combined our lifespan reference curves and harmonization mechanism into a Python package called eHarmonize. Dissemination of the package will allow collaborators to harmonize their data on-site and then share either the harmonized measures, or - in situations where raw data cannot be shared (e.g., genetics studies) - results of an agreed upon analysis. In addition, new sites will be free to join existing projects without requiring re-harmonization of the previously collected and harmonized data. We also created the package framework to be flexible and adaptable to update our existing reference curves and include reference curves for other measures, such as diffusivity measures or even those extracted from another modality. Version control ensures appropriate provenance for reproducibility.

The datasets we used to build our lifespan reference curves are public and therefore available to researchers for many applications. As such, researchers harmonizing multi-site data for machine learning studies may be concerned about data leakage with these datasets in the harmonization reference. After applying the eHarmonize framework to training and testing datasets, we found that there was no significant difference in MAE between training and testing datasets (p > 0.09). Our iterative sampling approach likely limited the overfitting of large datasets, such as the UK Biobank, so harmonization is not driven by any single dataset. eHarmonize may be an important tool for large scale machine learning purposes, but we recommend interested researchers to formally evaluate this for data leakage when using one of the training datasets, as was done in the establishment of *harmonizer*^*69*^.

Our lifespan reference curves have a few limitations that we plan to address in future versions. First, we had uneven sampling across the age range. In particular, children younger than 8 years old and adults in the range of 35 to 45 years old were underrepresented. We also acknowledge that the protocols of our training data were largely homogeneous, good quality data, and our test sets were similar. For the next iteration of lifespan curves, we aim to include more data from our underrepresented age ranges and acquisition protocols. To address the connection between acquisition protocol and biological interpretation between studies, we will also evaluate the added value of separate lifespan reference curves for different acquisition parameters.

To expand on eHarmonize’s functionality, future steps will include modeling functions that can take into account population-based sources of variability or additional model parameters. For the current study, we took care to only include unrelated and cross-sectional subsets of all studies. In the future, it would be important to examine nested random effects that account for a covariance (kinship) structure of the study population. There is currently one nested-ComBat package that implements both nested and Gaussian mixture model versions of ComBat^70^; however, the functions are currently hard-coded for bi-modal or binary data. With regard to model parameters, the World Health Organization (WHO) recommends GAM-LSS for creating lifespan charts^71^. GAM-LSS models may fit the first four moments of a distribution, adding in skew and kurtosis parameters, which GAM does not. Fitting more parameters robustly requires more data, and as many studies have small sample sizes, we elected to start with GAM models. For future studies wishing to combine only larger datasets, we will examine the effect of incorporating the ability to fit a GAM-LSS model. Additional features we will test include combining ComBat-GAM with CovBat to account for covariance between measures as well as the non-linear age trend.

Overall, we successfully created lifespan reference curves for regional FA measures and made progress in applying ComBat-GAM to these references. Our framework provides studies with the ability to standardize their dMRI measures. Collaborators from different sites can also harmonize their data to our lifespan reference curves without worrying about their covariate overlap or differences in sample size. The framework is now available as a Python package at https://github.com/ahzhu/eharmonize.

## Data Availability

All datasets used in building the lifespan reference and most of the test datasets are from publicly available datasets. For inquiries about the TAOS dataset, please contact Peter Kochunov. For inquiries about the GOBS dataset, please contact John Blangero and David Glahn.

## Code Availability

eHarmonize is an open-source package. The code may be read in full at https://github.com/ahzhu/eharmonize.

## Author Contributions

AHZ, TMN, PMT, and NJ conceptualized the study. AR, LS, GIdZ, KLM, MJW, SEM, JB, DCG, PK, and AKH provided access to study data. AHZ, TMN, and JEV-R processed images and aggregated data from the numerous datasets. AHZ created the lifespan reference curves, coded eHarmonize, and performed all statistical analysis. SJ helped with methods testing. AHZ, TMN, SJ and NJ wrote the manuscript.

## Acknowledgments

AHZ, TMN, SJ, JEV-R, PMT, and NJ received funding support from the NIH (R01MH134004, R01MH116147, R01AG058854, P41EB015922, RF1AG057892) and the Alzheimer’s Association. JB received funding support from the following NIH grants: U54HG013247, P30AG059305, U19AG076581, R01AG078423, and R01AG058464.

Data used in the preparation of this article were obtained from the Adolescent Brain Cognitive Development^SM^ (ABCD) Study (https://abcdstudy.org), held in the NIMH Data Archive (NDA). This is a multisite, longitudinal study designed to recruit more than 10,000 children age 9-10 and follow them over 10 years into early adulthood. The ABCD Study® is supported by the National Institutes of Health and additional federal partners under award numbers U01DA041048, U01DA050989, U01DA051016, U01DA041022, U01DA051018, U01DA051037, U01DA050987, U01DA041174, U01DA041106, U01DA041117, U01DA041028, U01DA041134, U01DA050988, U01DA051039, U01DA041156, U01DA041025, U01DA041120, U01DA051038, U01DA041148, U01DA041093, U01DA041089, U24DA041123, U24DA041147. A full list of supporters is available at https://abcdstudy.org/federal-partners.html. A listing of participating sites and a complete listing of the study investigators can be found at https://abcdstudy.org/consortium_members/. ABCD consortium investigators designed and implemented the study and/or provided data but did not necessarily participate in the analysis or writing of this report. This manuscript reflects the views of the authors and may not reflect the opinions or views of the NIH or ABCD consortium investigators. The ABCD data repository grows and changes over time. The ABCD data used in this report came from 10.15154/1523041. DOIs can be found at https://dx.doi.org/10.15154/1523041.

Data used in the preparation of this article were obtained from the Alzheimer’s Disease Neuroimaging Initiative (ADNI) database (adni.loni.usc.edu). The ADNI was launched in 2003 as a public-private partnership, led by Principal Investigator Michael W. Weiner, MD. The primary goal of ADNI has been to test whether serial magnetic resonance imaging (MRI), positron emission tomography (PET), other biological markers, and clinical and neuropsychological assessment can be combined to measure the progression of mild cognitive impairment (MCI) and early Alzheimer’s disease (AD). For up-to-date information, see www.adni-info.org.

Data collection and sharing for this project was funded by the Alzheimer’s Disease Neuroimaging Initiative (ADNI) (National Institutes of Health Grant U01 AG024904) and DOD ADNI (Department of Defense award number W81XWH-12-2-0012). ADNI is funded by the National Institute on Aging, the National Institute of Biomedical Imaging and Bioengineering, and through generous contributions from the following: AbbVie, Alzheimer’s Association; Alzheimer’s Drug Discovery Foundation; Araclon Biotech; BioClinica, Inc.; Biogen; Bristol-Myers Squibb Company; CereSpir, Inc.; Cogstate; Eisai Inc.; Elan Pharmaceuticals, Inc.; Eli Lilly and Company; EuroImmun; F. Hoffmann-La Roche Ltd and its affiliated company Genentech, Inc.; Fujirebio; GE Healthcare; IXICO Ltd.;Janssen Alzheimer Immunotherapy Research & Development, LLC.; Johnson & Johnson Pharmaceutical Research & Development LLC.; Lumosity; Lundbeck; Merck & Co., Inc.;Meso Scale Diagnostics, LLC.; NeuroRx Research; Neurotrack Technologies; Novartis Pharmaceuticals Corporation; Pfizer Inc.; Piramal Imaging; Servier; Takeda Pharmaceutical Company; and Transition Therapeutics. The Canadian Institutes of Health Research is providing funds to support ADNI clinical sites in Canada. Private sector contributions are facilitated by the Foundation for the National Institutes of Health (www.fnih.org). The grantee organization is the Northern California Institute for Research and Education, and the study is coordinated by the Alzheimer’s Therapeutic Research Institute at the University of Southern California. ADNI data are disseminated by the Laboratory for Neuro Imaging at the University of Southern California.

The Cambridge Centre for Ageing and Neuroscience (Cam-CAN) was supported by the UK Biotechnology and Biological Sciences Research Council (grant number BB/H008217/1), together with support from the UK Medical Research Council Cognition & Brain Sciences Unit (CBU) and University of Cambridge, UK. We are grateful to the Cam-CAN respondents and their primary care teams in Cambridge for their participation in the Cam-CAN study. We also thank colleagues at the MRC Cognition and Brain Sciences Unit MEG and MRI facilities for their assistance.

GOBS was supported by the National Institute of Mental Health MH0708143 (to D.C.G.), MH078111 (to J.B.), and MH083824 (to D.C.G. and J.B.).

Research reported in this publication was supported by the National Institute Of Mental Health of the National Institutes of Health under Award Number U01MH109589 and by funds provided by the McDonnell Center for Systems Neuroscience at Washington University in St. Louis. The HCP-Development 2.0 Release data used in this report came from DOI: 10.15154/1520708.

HCP data were provided [in part] by the Human Connectome Project, WU-Minn Consortium (Principal Investigators: David Van Essen and Kamil Ugurbil; 1U54MH091657) funded by the 16 NIH Institutes and Centers that support the NIH Blueprint for Neuroscience Research; and by the McDonnell Center for Systems Neuroscience at Washington University.

The Trøndelag Health Study (HUNT) is a collaboration between HUNT Research Centre (Faculty of Medicine and Health Sciences, Norwegian University of Science and Technology NTNU), Trøndelag County Council, Central Norway Regional Health Authority, and the Norwegian Institute of Public Health.

Data used in the preparation of this article were obtained from the Pediatric MRI Data Repository created by the NIH MRI Study of normal brain development. This is a multi-site, longitudinal study of typically developing children, from ages newborn through young adulthood, conducted by the Brain Development Cooperative Group and supported by the National Institute of Child Health and Human Development, the National Institute on Drug Abuse, the National Institute of Mental Health, and the National Institute ofNeurological Disorders and Stroke (Contract #s N01-HD02-3343, N01-MH9-0002, and N01-NS-9-2314, N01-NS-9-2315, N01-NS-9-2316, N01-NS-9-2317, N01-NS-9-2319 and N01-NS-9-2320). Disclaimer: The views herein do not necessarily represent the official views of the National Institute of Child Health and Human Development, the National Institute on Drug Abuse, the National Institute of Mental Health, the National Institute of Neurological Disorders and Stroke, the NIH, the US Department of Health and Human Services, or any other agency of the US Government.

Data were provided [in part] by OASIS-3: Longitudinal Multimodal Neuroimaging: Principal Investigators: T. Benzinger, D. Marcus, J. Morris; NIH P30 AG066444, P50 AG00561, P30 NS09857781, P01 AG026276, P01 AG003991, R01 AG043434, UL1 TR000448, R01 EB009352. AV-45 doses were provided by Avid Radiopharmaceuticals, a wholly owned subsidiary of Eli Lilly.

Data collection and subsequent dataset for this project were obtained from the Pediatric Imaging, Neurocognition and Genetics Study (PING), National Institutes of Health Grant RC2DA029475. PING is funded by the National Institute on Drug Abuse and the Eunice Kennedy Shriver National Institute of Child Health & Human Development. PING data are disseminated by the PING Coordinating Center at the Center for Human Development, University of California, San Diego, as detailed in Jernigan et al (2016).

Data used in the preparation of this article were obtained on August 25, 2022 from the Parkinson’s Progression Markers Initiative (PPMI) database (www.ppmi-info.org/access-data-specimens/download-data), RRID:SCR 006431. For up-to-date information on the study, visit www.ppmi-info.org. PPMI – a public-private partnership – is funded by the Michael J. Fox Foundation for Parkinson’s Research and funding partners, including 4D Pharma, Abbvie, AcureX, Allergan, Amathus Therapeutics, Aligning Science Across Parkinson’s, AskBio, Avid Radiopharmaceuticals, BIAL, BioArctic, Biogen, Biohaven, BioLegend, BlueRock Therapeutics, Bristol-Myers Squibb, Calico Labs, Capsida Biotherapeutics, Celgene, Cerevel Therapeutics, Coave Therapeutics, DaCapo Brainscience, Denali, Edmond J. Safra Foundation, Eli Lilly, Gain Therapeutics, GE HealthCare, Genentech, GSK, Golub Capital, Handl Therapeutics, Insitro, Janssen Neuroscience, Jazz Pharmaceuticals, Lundbeck, Merck, Meso Scale Discovery, Mission Therapeutics, Neurocrine Biosciences, Neuropore, Pfizer, Piramal, Prevail Therapeutics, Roche, Sanofi, Servier, Sun Pharma Advanced Research Company, Takeda, Teva, UCB, Vanqua Bio, Verily, Voyager Therapeutics, the Weston Family Foundation and Yumanity Therapeutics.”

The Queensland Twin IMaging (QTIM) study is forever grateful to the twins and siblings for their willingness to participate in our studies. We thank Marlene Grace and Ann Eldridge for participant recruitment; Kerrie McAloney for study co-ordination; Kori Johnson, Aaron Quiggle, Natalie Garden, Matthew Meredith, Peter Hobden, Kate Borg, Aiman Al Najjar and Anita Burns for data acquisition; David Butler and Daniel Park for IT support. The QTIM study was supported by the National Institute of Child Health and Human Development (R01 HD050735) and the National Health and Medical Research Council (496682, 1009064).

The Southwest University Longitudinal Imaging Multimodal (SLIM) Brain Data Repository was supported by the National Natural Science Foundation of China (31271087; 31470981; 31571137; 31500885), National Outstanding young people plan the Program for the Top Young Talents by Chongqing, the Fundamental Research Funds for the Central Universities (SWU1509383,SWU1509451), Natural Science Foundation of Chongqing (cstc2015jcyjA10106), Fok Ying Tung Education Foundation (151023), General Financial Grant from the China Postdoctoral Science Foundation (2015M572423, 2015M580767), Special Funds from the Chongqing Postdoctoral Science Foundation (Xm2015037), Key research for Humanities and social sciences of Ministry of Education(14JJD880009).

The TAOS study (PI: D.E. Williamson) was supported by the National Institute on Alcohol Abuse and Alcoholism (R01AA016274) —”Affective and Neuro-biological Predictors ofAdolescent-OnsetAUD” and the Dielmann Family.

This research has been conducted using data from UK Biobank, a major biomedical database, under application number 11559. UK Biobank is generously supported by its founding funders the Wellcome Trust and UK Medical Research Council, as well as the Department of Health, Scottish Government, the Northwest Regional Development Agency, British Heart Foundation and Cancer Research UK.

