## Supplementary Information for "Lifespan reference curves for harmonizing multi-site regional brain white matter metrics from diffusion MRI"

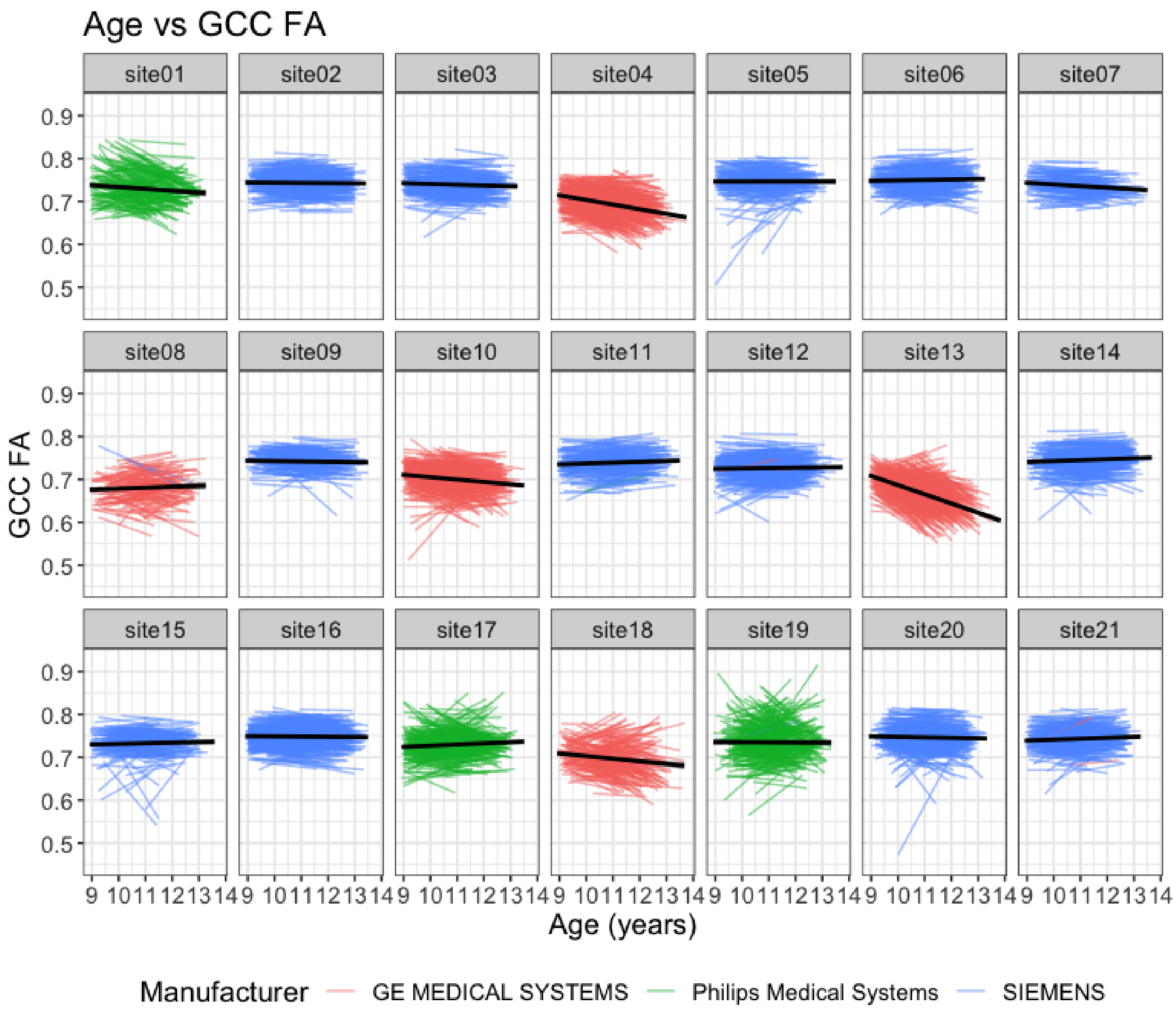

**Supplementary Figure 1.** ABCD site effects on FA in the genu of the corpus callosum (GCC). Longitudinal data from ABCD subjects are provided as spaghetti plots, separated by site and colored by manufacturer. The trend line per site is also included in black. Most sites with a GE scanner showed decreasing FA with age, contrary to the expected age effect. Philips was the least common scanner manufacturer. One Philips site had a downward trajectory (site01), and another had widely diverging subject-level trajectories (site19). As a result, we elected to use only ABCD data acquired on Siemens scanners.

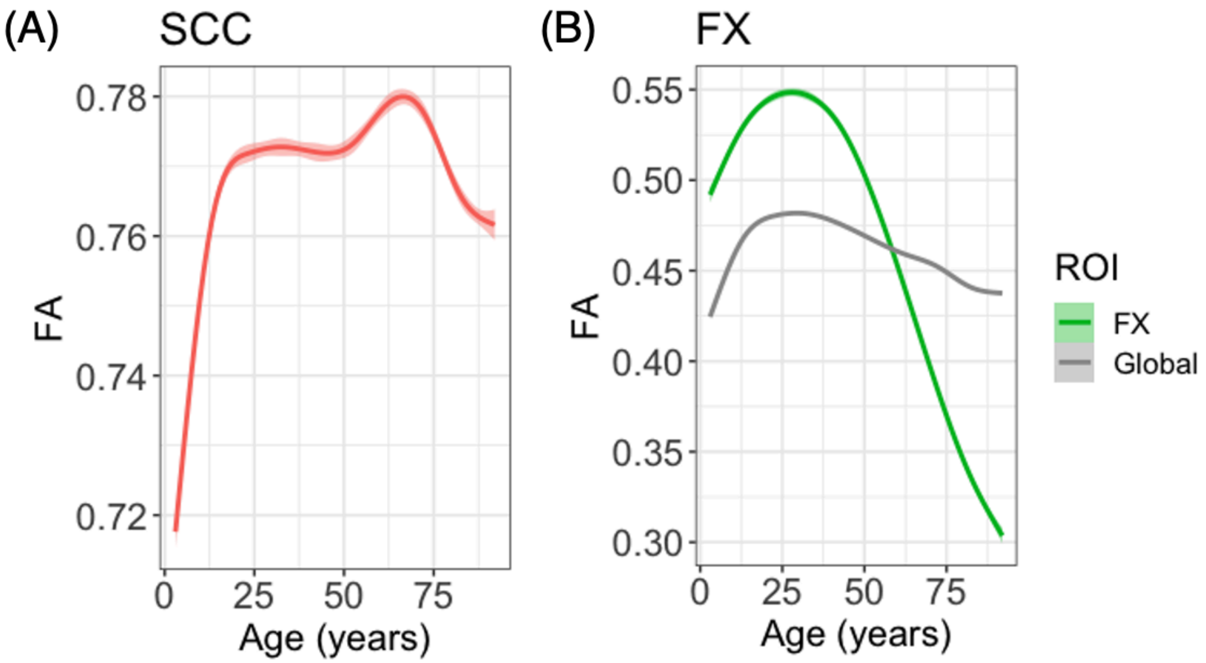

**Supplementary Figure 2.** Outlier ROI lifespan trajectories. (A) FA in the SCC peaks at age 68 years, much later than all other ROIs. (B) After peaking at age 29 years, FA in the FX decreases faster than in any other ROI. The lifespan reference curve for the global FA measure is also included for reference.

**Supplementary Table 1.** Sex effects across the white matter. ROIs where males had significantly higher FA are in **bold**. ROIs where females had significantly higher FA are *italicized*.
* $p < 0.05$; ** $p < 0.001$

| ROI | Estimate | SE | t-value | p-value | pFDR |
|---|---|---|---|---|---|
| **AverageFA** | **0.0020** | **0.00024** | **8.4** | **< 0.001**** | **< 0.001**** |
| **ACR** | **0.0018** | **0.00037** | **4.8** | **< 0.001**** | **< 0.001**** |
| **ALIC** | **0.0042** | **0.00040** | **10.5** | **< 0.001**** | **< 0.001**** |
| BCC | -0.00078 | 0.00046 | -1.7 | 0.089 | 0.092 |
| CC | 0.00014 | 0.00038 | 0.4 | 0.71 | 0.71 |
| **CGC** | **0.010** | **0.00048** | **21.7** | **< 0.001**** | **< 0.001**** |
| **CGH** | **0.0049** | **0.00066** | **7.4** | **< 0.001**** | **< 0.001**** |
| **CR** | **0.0013** | **0.00031** | **4.1** | **< 0.001**** | **< 0.001**** |
| **CST** | **0.0082** | **0.00053** | **15.5** | **< 0.001**** | **< 0.001**** |
| **EC** | **0.0030** | **0.00034** | **8.7** | **< 0.001**** | **< 0.001**** |
| *FX* | *-0.0068* | *0.00082* | *-8.3* | *< 0.001*** | *< 0.001*** |
| *FXST* | *-0.0050* | *0.00047* | *-10.5* | *< 0.001*** | *< 0.001*** |
| **GCC** | **0.0012** | **0.00048** | **2.5** | **0.011*** | **0.013*** |
| **IC** | **0.0032** | **0.00032** | **10.0** | **< 0.001**** | **< 0.001**** |
| *PCR* | *-0.00077* | *0.00037* | *-2.1* | *0.040* | *0.046* |
| **PLIC** | **0.0041** | **0.00036** | **11.5** | **< 0.001**** | **< 0.001**** |
| *PTR* | *-0.0035* | *0.00044* | *-7.9* | *< 0.001*** | *< 0.001*** |
| **RLIC** | **0.00082** | **0.00039** | **2.1** | **0.036** | **0.043** |
| SCC | 0.00076 | 0.00041 | 1.9 | 0.063 | 0.069 |
| **SCR** | **0.0014** | **0.00037** | **3.9** | **< 0.001**** | **< 0.001**** |
| **SFO** | **0.0044** | **0.00042** | **10.6** | **< 0.001**** | **< 0.001**** |
| **SLF** | **0.0030** | **0.00035** | **8.6** | **< 0.001**** | **< 0.001**** |
| *SS* | *-0.0051* | *0.00043* | *-11.9* | *< 0.001*** | *< 0.001*** |
| *TAP* | *-0.0034* | *0.00069* | *-4.9* | *< 0.001*** | *< 0.001*** |
| **UNC** | **0.0050** | **0.00060** | **8.4** | **< 0.001**** | **< 0.001**** |

**Supplementary Table 2.** ApoE4 effects across the white matter. ROIs where ApoE4 was associated with significantly lower FA after multiple comparisons correction are in **bold**.
* p < 0.05; ** p < 0.001

| ROI | N | Estimates | SE | t_values | p_values | p_FDR |
|---|---|---|---|---|---|---|
| **AverageFA** | **26902** | **-0.020** | **0.0057** | **-3.42** | **< 0.001**** | **0.0052*** |
| ACR | 26902 | -0.0081 | 0.0052 | -1.56 | 0.12 | 0.25 |
| ALIC | 26902 | -0.0039 | 0.0059 | -0.67 | 0.50 | 0.63 |
| BCC | 26902 | -0.0084 | 0.0060 | -1.39 | 0.16 | 0.27 |
| CC | 26902 | -0.014 | 0.0060 | -2.27 | 0.02* | 0.097 |
| CGC | 26902 | -0.010 | 0.0058 | -1.76 | 0.08 | 0.20 |
| **CGH** | **26902** | **-0.027** | **0.0059** | **-4.61** | **< 0.001**** | **< 0.001**** |
| CR | 26902 | -0.0045 | 0.0056 | -0.80 | 0.42 | 0.56 |
| CST | 26902 | -0.0054 | 0.0059 | -0.92 | 0.36 | 0.50 |
| EC | 26902 | -0.0013 | 0.0059 | -0.22 | 0.83 | 0.90 |
| FX | 26902 | -0.0063 | 0.0044 | -1.43 | 0.15 | 0.27 |
| FXST | 26902 | -0.011 | 0.0056 | -1.98 | 0.048* | 0.13 |
| GCC | 26902 | -0.012 | 0.0056 | -2.10 | 0.036* | 0.13 |
| IC | 26902 | -0.0060 | 0.0060 | -1.01 | 0.31 | 0.46 |
| PCR | 26902 | -0.0078 | 0.0060 | -1.32 | 0.19 | 0.29 |
| PLIC | 26902 | 0.00043 | 0.0060 | 0.071 | 0.94 | 0.94 |
| **PTR** | **26902** | **-0.022** | **0.0057** | **-3.78** | **< 0.001**** | **0.0020*** |
| RLIC | 26902 | -0.012 | 0.0061 | -2.05 | 0.040* | 0.13 |
| **SCC** | **26902** | **-0.016** | **0.0059** | **-2.70** | **0.0068*** | **0.043*** |
| SCR | 26902 | 0.0031 | 0.0060 | 0.51 | 0.61 | 0.69 |
| SFO | 26902 | 0.00081 | 0.0060 | 0.14 | 0.89 | 0.93 |
| SLF | 26902 | -0.0091 | 0.0060 | -1.52 | 0.13 | 0.25 |
| SS | 26902 | -0.015 | 0.0060 | -2.56 | 0.01* | 0.052 |
| TAP | 26902 | -0.0096 | 0.0060 | -1.59 | 0.11 | 0.25 |
| UNC | 26902 | -0.0033 | 0.0060 | -0.54 | 0.59 | 0.69 |
